# Burn intensity regulates post-fire effects on soil N_2_O emissions: A grassland soil pyrocosm study

**DOI:** 10.64898/2026.10.27.761429

**Authors:** Kenneth M Czapla, Elizah Z Stephens, Alexander H Krichels, Maria E Ordoñez, Basubi B Zhilik, Luke S Hillary, Steven D Allison, Michael J Wilkins, Joanne B Emerson, Sydney I Glassman, Peter M Homyak

## Abstract

Post-fire soil emissions of nitric oxide (NO) and nitrous oxide (N_2_O) are important ecosystem nitrogen (N) loss pathways, but their flux dynamics vary widely across studies, making generalizations challenging. Natural variation in burn intensity—or heat energy released during combustion—may help resolve this variability by mediating the soil physicochemical and biological drivers of these emissions. Here, we used an experimental "pyrocosm" system to evaluate how varying burn intensity influences post-fire NO and N_2_O emissions using soil from a Southern California grassland across 18 weeks. Over several months, we measured soil NO and N_2_O fluxes alongside soil pH, extractable nitrate (NO_3_^-^) and ammonium (NH_4_^+^), net NH_4_^+^ production, net nitrification, soil carbon (C) pools, microbial biomass C, and the potential activities of five extracellular hydrolytic enzymes under Control, Low-(100.1 ± 20.0 °C peak soil temperature at 5cm), and High-intensity (243.8 ± 7.4 °C peak soil temperature at 5cm) burns. Soil NO emissions were not significantly affected by burn intensity. In contrast, N_2_O emissions tripled under low burn intensity relative to unburned controls, but did not differ from controls at high burn intensity. This stimulation of N_2_O emission under low burn intensity was driven by increased denitrification, as supported by elevated soil pH and labile C availability, and an 88% reduction in net nitrification that signaled active NO_3_^-^consumption. Under high burn intensity, however, severe loss of microbial biomass, soil C, and NO_3_^-^constrained denitrification, with rising hydrolytic enzyme activities over time suggesting C limitation. Together, these findings demonstrate that post-fire N_2_O emissions reflect a fire intensity-dependent balance between microbial mortality and resource limitation, helping to explain divergent N emissions across ecosystems.

## 1. Introduction

As global wildfires become more frequent and intense (Cunningham et al., 2024), understanding post-fire ecosystem nitrogen (N) loss is necessary for predicting changes in productivity, as N is often lost during wildfires but is essential for ecosystem recovery (Hanan et al., 2016). While wildfires directly volatilize N during biomass combustion, significant N loss may continue long after fires are extinguished through post-fire leaching and soil emission of trace gases, including nitric oxide (NO; an air pollutant at high concentrations) and nitrous oxide (N_2_O; a greenhouse gas ∼300 times more powerful than carbon dioxide; Mannucci et al., 2015; Yoro et al., 2020; Lindaas et al., 2021). Post-fire soil N emissions vary greatly within and between ecosystems, but the factors regulating this variability remain unresolved (Stephens & Homyak, 2023). Variation in burn intensity—the rate of energy released as heat during combustion (Keely 2009)—can dictate the severity of soil physicochemical alterations and microbial mortality that govern microbial NO and N_2_O emission production (Holden & Treseder, 2013; Stephens et al., 2026) and, thus, may help explain post-fire N gas emission variability.

Although fire intensity varies widely within and across landscapes due to differences in fuel load, wind speeds, temperatures, humidity, soil and vegetation moisture, and vegetation composition (Davies et al., 2016; Haas et al., 2022), it is rarely measured directly during active wildfires. Therefore, evaluating whether variation in burn intensity explains post-fire NO and N_2_O emissions is essential to improving our ability to predict, scale up, and manage ecosystem fire impacts.

The net effect of burn intensity on post-fire NO and N_2_O emissions likely reflects the balance between at least two competing soil mechanisms: 1) changes in soil physicochemical factors such as increased pH and availability of N substrates that stimulate N gas emissions and 2) microbial mortality and alteration of the biological processes that emit NO and N_2_O. Higher burn intensities may stimulate emissions by increasing both soil pH and ammonium (NH_4_^+^) availability (Certini et al., 2005; Stephens et al., 2026) as greater burn intensity deposits more ash, raising soil pH (Goforth et al. 2005), while thermal mineralization of soil organic matter increases NH_4_^+^ availability. In turn, elevated soil pH and NH_4_^+^ availability favor both nitrification—the aerobic oxidation of ammonia (NH_3_) to nitrate (NO_3_^-^)—and denitrification— the anaerobic reduction of NO_3_^-^ to N_2_, both of which release NO and N_2_O as byproducts (Firestone & Davidson 1989; Niu et al., 2024). Importantly, environmental factors such as soil moisture regulate the net balance between NO production and consumption; under dry conditions, NO can diffuse out of the soil matrix, whereas elevated moisture restricts gas diffusivity, favoring anaerobic processes, microbial NO consumption, and its subsequent reduction to N_2_O (Schindlbacher et al., 2004). Conversely, high intensity fires can simultaneously suppress microbial processes (Holden et al., 2016; Su et al., 2022) and drive significant microbial mortality (Holden & Treseder, 2013; Pressler et al., 2019; Whitman et al., 2019; Pulido-Chavez et al., 2023). Furthermore, high intensity fire can deplete labile organic C pools (Gonzalez-Perez et al., 2004; Krichels et al., 2025) and inhibit extracellular C- and N-acquiring enzyme activity (Jones et al., 2010), limiting acquisition of C and N required for denitrification. Because burn intensity affects the balance between these stimulating and inhibiting mechanisms, post-fire NO and N_2_O emissions may vary with burn intensity.

Grasslands represent a critical biome for testing the role of burn intensity in regulating N emissions. Grassland ecosystems experience roughly 80% of global wildfire events (Leys et al., 2018) and account for approximately 18% of global NO emissions (Tian et al., 2023) and 30% of global N_2_O emissions (Yu & Zhuang, 2019). Furthermore, grassland fires—ranging from prescribed burns to wildfires—have wide spatial variation in burn intensity depending on fuel loads, fuel moisture, and microclimatic conditions (Davies et al., 2016; Stavi et al., 2019; Fusco et al., 2022). Thus, resolving how burn intensity controls grassland N gas fluxes is essential for scaling regional and global NO and N_2_O budgets. To understand whether burn intensity resolves variability in grassland N gas losses, we used soil from a grassland where we had previously studied effects of prescribed and wildfire burning on soil chemistry, microbiome, and vegetation responses (Glassman et al., 2023) to ask: 1) How does varying burn intensity influence post-fire soil NO and N_2_O emissions? and 2) How does burn intensity impact the key physical, chemical, and biological soil drivers of these emissions (pH, NH_4_^+^, NO_3_^-^, net N mineralization/nitrification, C availability, microbial biomass, and extracellular enzyme activities)?

To address these questions, we used a controlled “pyrocosm” approach, burning experimental mesocosms of homogenized, sieved soil (Bruns et al., 2020) from which we subsampled soils over 18 weeks. By minimizing natural soil heterogeneity, this controlled pyrocosm framework isolated the direct effects of burn intensity on trace gas emissions with high burn temperature reproducibility. We evaluated two burn treatments: a low-intensity burn aimed to approximate the heat exposure characteristic of natural grassland fires, which are typically fast-moving, fine-fuel-driven events that drive limited thermal penetration into the soil profile (Stavi 2019); and an extreme high-intensity burn intended to exceed natural grassland intensity. By contrasting a realistic fire scenario against an extreme burn intensity, we aimed to isolate the specific biological (e.g., microbial biomass and activity) and geochemical (e.g., fire-generated N pools) drivers underlying gaseous N fluxes, and to clarify which of these are most sensitive (or resistant) to fire-driven disturbance. To capture post-fire trace gas pulses, soil subsamples from the pyrocosms were periodically wetted to 80% water-holding capacity during microcosm flux assays, approximating post-fire rainfall events followed by soil drying. We hypothesized that increasing burn intensity would elevate denitrifier- and nitrifier-derived NO and N_2_O emissions by raising soil pH, NH_4_^+^ availability, and microbial access to C measured as CO_2_ respired during incubations (hereafter referred to as immediately respirable C; IRC). Conversely, we hypothesized that extreme burn intensities would limit or suppress these emissions due to severe declines in microbial biomass, labile C availability, and extracellular enzyme activities.

## 2. Methods

### 2.1. Site description

We collected soil from a *Bromus*-dominated grassland in the Santa Rosa Plateau Ecological Reserve (SRPER; 560 m elevation; Figure S1) in Riverside County, CA, USA. SRPER has a Mediterranean climate typical of Southern California, with annual rainfall averaging about 762 mm (occurring mostly from November to April), and average monthly temperatures lowest during December at 11.5 °C and peaking during August at 24.1 °C (weathercurrents.com/murrieta/Climate.do). Parts of SRPER burned in 2018 (prescribed) and in 2019 (Tenaja wildfire) and post-fire impacts to biogeochemical characteristics, plants, and microbiomes were previously characterized (Glassman et al., 2023). For our study, soil from the top 15 cm was collected with shovels on May 21, 2024, within 10 meters of 33°32’02.0"N 117°15’15.9"W, located in an unburned area of SRPER adjacent to unburned plots of the Tenaja wildfire study (Glassman et al., 2023). Soils at the site have a rocky loam texture classified as Lithic Ruptic-Inceptic Haploxeralfs mapped within the Vallecitos series (https://websoilsurvey.nrcs.usda.gov).

### 2.2. Experimental design

Soil “pyrocosms” were burned on July 10, 2024. Pyrocosms were created by filling 9 L galvanized steel buckets 15 cm deep with air-dried soils that were homogenized and sieved to 2mm (Bruns et al., 2020, Figure S2). We placed thermocouples (K Type Thermocouple Probe 1m -50-800 Degree Centigrade Mini Connector Temperature Sensors, E-outstanding, Shenzhen, China) at 5 and 10 cm soil depths on each pyrocosm to record temperature during the burn until temperature stabilized to ambient at around 24 h with dataloggers (Sper Scientific Certified Bluetooth 4 Channel Datalogging, Scottsdale, AZ, USA). Each pyrocosm was then randomly assigned to one of three treatments: “Low burn,” “High burn,” and unburned “Control” (n = 8 per treatment). Different masses of all-natural charcoal (B&B oak briquettes) were burned to manipulate burn intensity (100 g for the Low and 320 g for the High treatment). To increase the mass of soil sample available for biogeochemical measurements, pyrocosms were paired according to similarity in peak burn temperature and one core from each pyrocosm in a pair was composited into a single replicate, thus n = 4 per treatment.

To facilitate ignition, we built cardboard cylindrical structures held together with wooden dowels containing 40 g dried grass kindling gathered from the site and crumpled newspaper underneath and charcoal briquettes resting on top of the structure for each pyrocosm (Bruns et al 2020). The entire structure was burned until the charcoal briquettes rested on the soil surface and ashed completely overnight. Control pyrocosms were not burned, but were placed outdoors on a cement platform at the UCR Agricultural Operations Station alongside the two burn treatments during the experimental burn. Once burned, all pyrocosms were allowed to cool outdoors overnight and then moved to the laboratory to incubate at room temperature (∼22 °C) in four randomly arranged blocks for the duration of the 18-week experiment. Pyrocosms were covered with aluminum foil throughout the experiment to minimize cross-contamination and aerial microbial dispersal.

We began sampling pyrocosm soils one week after burning. One soil core 10 cm deep and 2.7 cm in diameter was taken from each pyrocosm in a pair, using sterile 60 mL syringes with cut-off ends, and paired cores were homogenized. Soil water holding capacity (WHC) was measured one week after burning by saturating 15 g soil from each pyrocosm with water, allowing water to drain against gravity through a 10 µm pore size paper filter situated in a funnel for 16 hours in a sealed container to prevent evaporation, and measuring moisture content to represent 100% WHC. Then, we watered the pyrocosms using ice (511 g, equating to ∼8 mm rainfall) that thawed overnight to slowly increase the soil water content from 7.4 to 20–50 % WHC (Figure S3b) and stimulate microbial activity in the pyrocosm. This sampling process was repeated at 3, 5, 9, 13, and 18 weeks after burning, with ice being added every two weeks (ice mass was proportionally reduced from first ice addition to account for decreasing soil surface area remaining after each coring) immediately after soil samples were collected from the pyrocosms until week 13 (i.e., soil subsamples were collected from pyrocosms at their driest point— < 20% WHC—just before raising soil moisture again with ice). To keep the soils from becoming too wet, we reduced the amount of ice to 250 g for the final pyrocosm wetting to not exceed 50% WHC. To parallel the pyrocosm wetting scheme, soil subsamples from pyrocosms were incubated in mason jars and wet to 80% WHC each day for 3 days and then allowed to dry to ∼30% for four days during trace gas flux analysis (Figure S4b).

We measured variables for each timepoint from three distinct points during the experimental procedure for each sampling: 1) soil pH, WHC, soil extractable NH_4_^+^ and NO_3_^-^, and total soil %C directly from the pyrocosm soil cores wetted every two weeks with ice, 2) NO and N_2_O fluxes, net NH_4_^+^ production and net nitrification, and immediately respirable carbon (IRC) during 7 day incubations in mason jars using soil subsamples obtained from the pyrocosm cores wetted to 80% WHC, and 3) microbial biomass C and potential enzyme activities from mason jar soil subsamples after they were dried to below 30% WHC at the end of the incubations (Figure S4).

### 2.3. Soil physicochemical measurements

To measure pyrocosm soil moisture at each time point, a ∼10 g subsample was weighed before and after oven drying at 105 °C for 24 h. pH was also measured at each time point with a calibrated Orion VersaStar Pro Benchtop pH Meter using 10 g dry weight (dw) soil shaken for 30 seconds with 20 mL deionized water (> 17.8 mΩ) and measured immediately.

Dried soils were ground and weighed to 30 mg in tin capsules and analyzed for total %C and %N content on an elemental analyzer (Flash EA 1112 CN; Thermo Fisher Scientific Inc). Soil extractable inorganic N was extracted from a 3 g soil subsample with 30 mL 2 M KCl solution that was shaken for 30 minutes and gravimetrically filtered with Whatman 42 filters on the day of sampling. KCl extracts were refrigerated overnight and analyzed for NO_3_ ^-^ (SEAL method EPA-129-A) and NH_4_^+^ (SEAL method EPA-126-A) on an AQ2 discrete autoanalyzer (SEAL Analytical Inc, Milwaukee, WI, USA) the following day.

### 2.4. Soil NO, N_2_O, and C Flux Measurements

NO, N_2_O, and CO_2_ fluxes were measured on a continuous flowthrough gas analysis system linking a Scintrex LMA-3 chemiluminescent NO_2_ analyzer, laser spectroscopy N_2_O analyzer (Model 914-0027, Los Gatos Research Inc., Mountain View, CA), and a LI-COR 8100 (LI-COR Biosciences, Lincoln, NE, USA; Krichels et al., 2024; Zhao et al., 2025). For each sample, 50 g soil was placed in a 4 oz glass mason jar and wet to 80% WHC 15 minutes before the first flux measurement, and a LI-COR 8150 multiplexer was used to measure fluxes in each of 12 jars once every 2.5 h for 7 days. To detect N_2_O produced by nitrification, ^15^NH_4_^+^ tracer was added at 2 atom % of soil NH_4_ ^+^ (see Supplementary Methods for equations used to calculate nitrification production of N_2_O from tracer). Because gas sample flowing at 1.5 L min^-1^ through the loop system caused drying at a rate of ∼15-20 % WHC per day, each sample was rewet to 80% WHC 24 and 48 h after the initial wetting, and then allowed to dry to below 30% WHC for the remainder of the seven-day incubation. This wetting scheme paralleled the wetting of pyrocosms with ice every two weeks and slowly air-drying between samplings. Before flowing through the instruments, water vapor in the headspace was removed by flowing through Nafion (PD-200 T-12MPS, Perma Pure LLC, Lakewood Township, NJ, USA) to avoid interference of moisture with gas measurements. The NO_2_ analyzer was equipped with a CrO_3_ NO oxidizer (Drummond Technology Inc., Ontario, Canada) to convert NO to NO_2_ and calibrated at the beginning and end of measurements using an eight-point standard curve (120 ppb NO, Airgas) diluted with zero grade air (Airgas). We spot-checked the gas measurements periodically by bypassing the CrO_3_ oxidizer to confirm that our soil was primarily producing NO; NO_2_ was not detectable, as expected. Because chemiluminescent analysis consumes NO within the gas sample, the NO_2_ analyzer was split off from the sample loop using timed 3-way valves and analyzed for the first 5 minutes so that it did not recycle gas back into the soil jars, and then for the final 5 minutes the valves switched back to recycling the gas sample to the jars to measure N_2_O and CO_2_. Flux jar headspace air flowing through NO_2_ analyzer at 1.5 L min^-1^ was replaced with zero grade air.

Fluxes were calculated using a custom R script (Andrews & Krichels, 2022), taking the slopes of the final 200 seconds for N_2_O and CO_2_ fluxes. Slopes of fluxes were typically highly linear (R^2^ > 0.98). For NO flux calculations, the final 30 seconds was used, and a mixing model with zero grade air was applied to calculate fluxes from equilibrated NO concentrations:

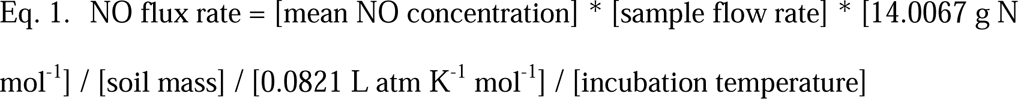

Cumulative fluxes for each timepoint were calculated using the “Trapz” function in R (pracma package, Borchers 2025), and these cumulative fluxes were summed to calculate total fluxes. Thus, we did not attempt to extrapolate gas fluxes over large gaps between measurements, and the total fluxes are only representing the time soils were measured (6 out of the 18 total weeks of the experiment). Cumulative CO_2_ fluxes integrated over 7 days were used as an index of C immediately available to microbes, hereafter defined as immediately respirable C (IRC; Homyak et al., 2018).

Immediately after flux measurements, incubation jars were subsampled for KCl-extracted NH_4_ ^+^ and NO_3_ ^-^, soil microbial biomass, and potential extracellular enzyme activity. NH_4_ ^+^, NO_3_ ^-^, and microbial biomass C subsamples were extracted, and subsamples taken to measure extracellular enzyme activities were stored for up to 11 months at -20 °C until analysis. We calculated net NH_4_ ^+^ production as the difference between KCl-extracted NH_4_^+^ determined before and after the 7-day flux measurements, and net nitrification as the difference between KCl-extracted NO ^-^ before and after the flux measurements.

### 2.5. Microbial biomass and potential extracellular enzyme activities

The soil subsamples for microbial biomass taken after flux incubations from the mason jars were fumigated by shaking 10 g soil for 4 hours with 40 mL 0.05 M K_2_SO4 and 0.5 mL chloroform (Fierer & Schimel, 2002). These samples and identical sets of samples without chloroform addition were shaken, bubbled for 20 minutes with lab air to remove residual chloroform, filtered with a Whatman 42 ashless cellulose filter, and analyzed for total organic C (TOC) on a Shimadzu TOC-V analyzer. Microbial biomass C was calculated as the difference in TOC between chloroform-fumigated and unfumigated subsamples, and because extraction efficiency was not corrected for this represents a flush of microbial C rather than biomass.

Extracellular enzyme activity (EEA) assays were performed as described in Püspök et al. (2022). Briefly, three extracellular hydrolytic C degradation enzymes α-glucosidase (AG), β-glucosidase (BG), 1,4-β-cellobiohydrolase (CBH), a hydrolytic P acquisition enzyme phosphomonoesterase (PHO), and a hydrolytic N acquisition enzyme N-acetyl-β-d-glucosaminidase (NAG) were assessed for the three pyrocosm treatments at the six time points over the course of the experiment. Enzyme potential activities were traced with the following fluorescing methylum-belliferone (MUB) or 7-amido-4-methyl-coumarin (MC)-tagged enzyme substrates: AG with 4-MUB-α-d-glucopyranoside, BG with 4-MUB-β-d-glucopyranoside, CBH with 4-MUB-β-d-cellobioside, PHO with 4-MUB-phosphate, and NAG with 4-MUB-N-acetyl-β-d-glucosaminide (Marx et al., 2001). Assays were performed on 96-well plates with 100 µL soil slurry mixed with 100 µL substrate solution, and after incubating the assays for 24 hours, fluorescence was read at 460 nm with a Tecan plate reader five times over 3 hours (Saiya-Cork et al. 2002, McLaren et al. 2017).

### 2.7. Statistics

R (version 4.5.2, R Core Team, 2024) was used to test for significant differences between treatments with one-way ANOVA tests (α = 0.05) in cases such as peak burn temperatures or cumulative fluxes where time is not a factor. ANOVA assumptions of normality were checked using histograms, q-q plots, and Shapiro-Wilk tests, and no transformations were needed to achieve data normality except for cubic root transformations on N_2_O measurements. To compare averages across treatments and time points, linear mixed-effect models (LMEs) were constructed using the “lme4” package (Bates et al., 2015), and the “emmeans” package was used to perform Tukey post-hoc tests with an adjustment for repeated measures (Lenth et al., 2026). Linear regressions were used to detect correlations between peak burn temperatures and dependent variables. Statistical results are summarized in Table S1.

## 3. Results

### 3.1 Burn Temperatures

The peak burn temperature measured at 5cm depth in Low burn pyrocosms averaged (hereafter, ± standard error) 100.1 ± 20.0 °C and was significantly lower than the peak temperature in High burn pyrocosms (243.8 ± 7.4°C; p < 0.017, ANOVA, Figure S5). The burn treatments also had highly replicable temperature profiles over time; they reached peak temperatures ∼5 hours after ignition for the Low burn and ∼10 hours for the High burn, with temperatures descending back to pre-burn temperatures after ∼21 hours for both burn treatments. Because the burn temperature profiles were consistent across replicates, we used peak burn temperatures at 5cm depth as a proxy for burn intensity.

### 3.2 Soil physicochemical measurements

Soil pH averaged 6.25 ± 0.03 before the burn. After the burn, soil pH increased to 7.27 ± 0.29 in the High burn and to 6.63 ± 0.03 in the Low burn (Figure 1a; p < 0.0001 for High vs. Control and p = 0.002 for Low vs. Control, LME). High burn soil pH remained higher than the Low burn throughout the 18-week experiment (p < 0.0001, LME).

**Figure 1.**
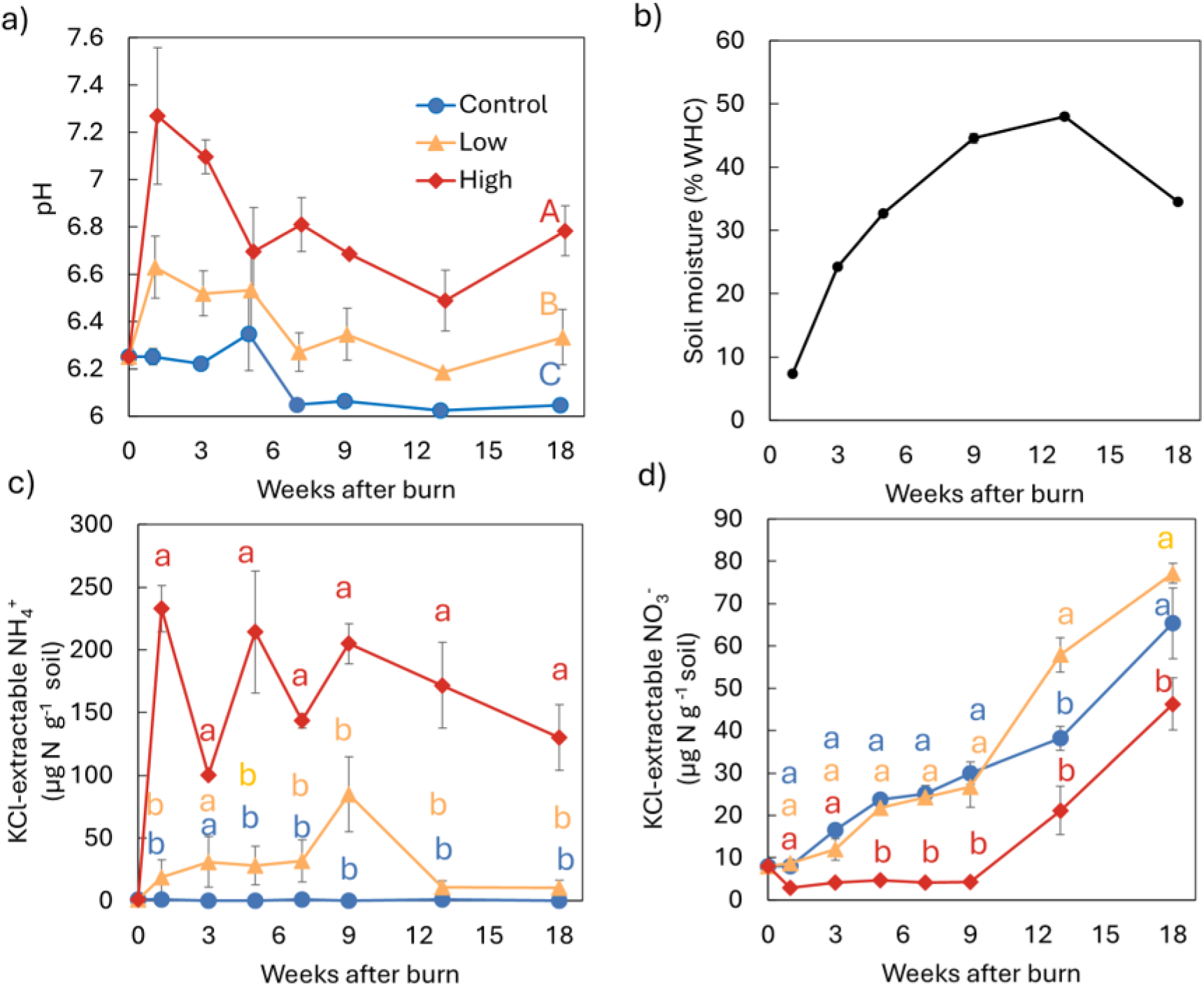
Dynamics of soil physicochemical properties across burn intensity treatments over an 18-week incubation. Panels display (a) soil pH, (b) soil moisture content expressed as percent water-holding capacity (WHC; averaged across all treatments), (c) KCl-extractable NH_4_^+^, and (d) KCl-extractable NO_3_^-^. Pyrocosm treatments include unburned Control, Low burn intensity (peak temperature of 100 ± 20 °C at 5 cm depth), and High burn intensity (244 ± 7 °C at 5 cm depth). Different letters denote statistically significant differences (p < 0.05) based on linear mixed-effect models (lowercase letters indicate treatment differences within individual time points; uppercase letters represent main treatment effects). Overlapping points were jittered for error bar visibility. Error bars represent standard errors (n = 4).

At 100% WHC, soils averaged 0.54 ± 0.01 g H_2_O g^-1^ soil in both the Control and Low burn, but this was reduced to 0.46 ± 0.01 g H_2_O g^-1^ soil in the High burn (p < 0.01, ANOVA; Figure S3). As we wetted pyrocosm soils during our experiment, soil moisture in the top 10 cm increased gradually from 7.4% to a maximum of 48% WHC at week 13, decreasing to 34.5% WHC by the end of the experiment (Figure 1b).

Burning increased soil extractable NH_4_^+^ by 17% in the Low burn and by 214% in the High burn relative to the Control (p < 0.001, LME; Figure 1c). In contrast to NH_4_^+^, the Low burn had no effect on extractable NO_3_^-^ (p = 0.7, LME); however, the High burn decreased extractable NO_3_ ^-^ by 2.7 times relative to the control (p < 0.0001, LME; Figure 1d). For all treatments, NO_3_^-^ was significantly greater than the previous timepoint starting at week 9, indicating a trend of increasing NO_3_^-^ over weeks 9-18 regardless of burn treatment (p < 0.01, LME).

### 3.3 Soil N Cycling Measurements

Soil NO emissions (Figure 2a,b) were not significantly impacted by burn intensity (p = 0.27, LME). Total cumulative NO emissions over the experiment averaged 6.13 ± 0.14 µmol N-NO g^-1^ for Control, 7.15 ± 0.69 µmol N-NO g^-1^ for Low burn, and 9.02 ± 0.88 µg N-NO g^-1^ for High burn. Although NO emissions began to increase in the High burn compared to Low and Control treatments 18 weeks after the burn, neither cumulative emissions nor individual time-point emissions differed significantly among Control, Low burn, and High burn treatments (p > 0.27, LME).

**Figure 2.**
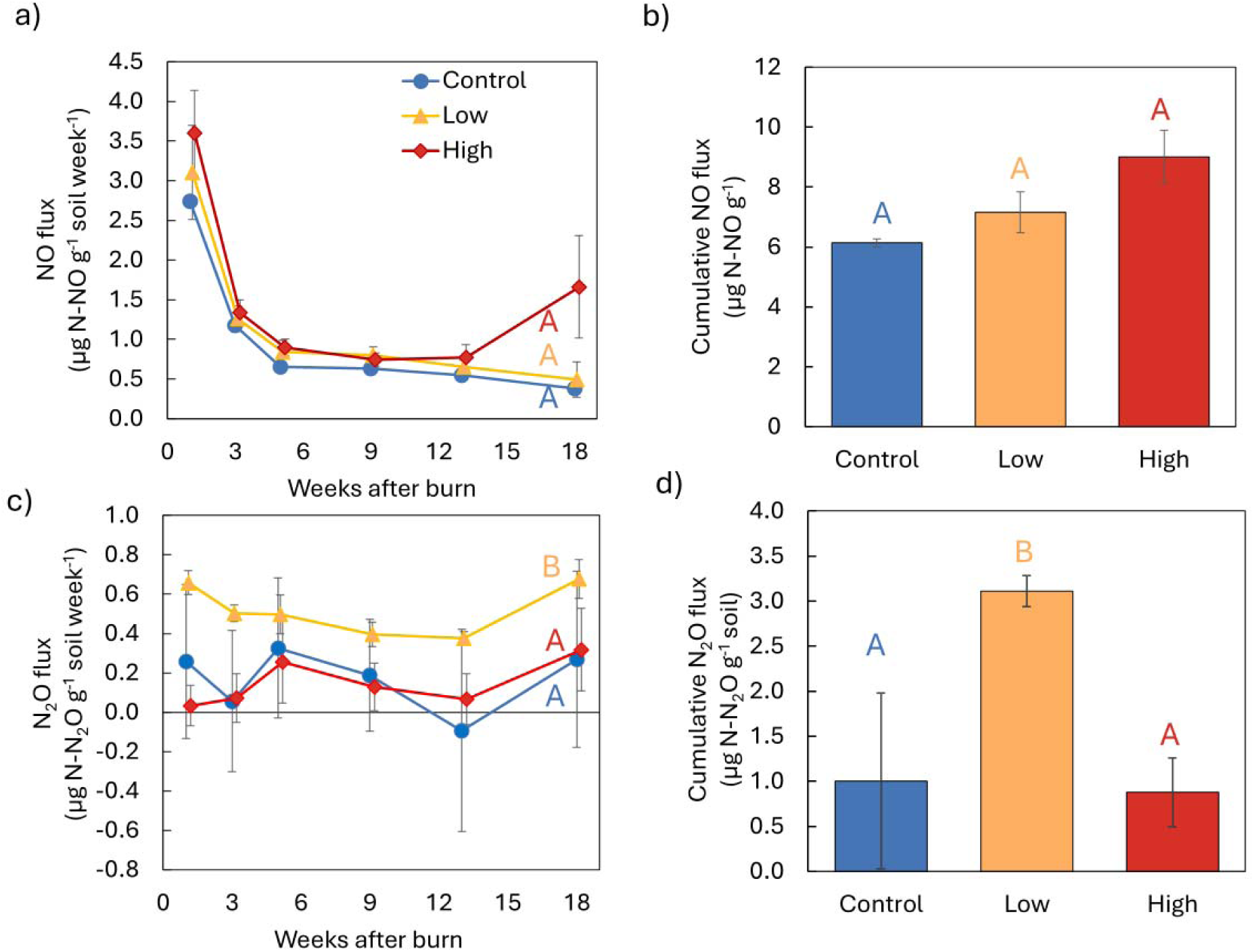
Soil trace nitrogen gas emissions across burn intensity treatments over an 18-week incubation. Panels show time-series flux rates for (a) NO and (c) N_2_O, alongside cumulative fluxes for (b) NO and (d) N_2_O integrated over the 18-week incubation. Pyrocosm treatments include unburned Control, Low burn intensity (peak temperature of 100 ± 20 °C at 5 cm depth), and High burn intensity (244 ± 7 °C at 5 cm depth). Different letters denote statistically significant differences (p < 0.05) based on linear mixed-effect models for time-series rates (a, c) and one-way ANOVAs for cumulative fluxes (b, d; lowercase letters indicate treatment differences within individual time points; uppercase letters represent main treatment effects). Overlapping points were jittered for error bar visibility. Error bars represent standard errors (n = 4).

In contrast to NO, low burn intensity stimulated a three-fold increase in N_2_O emissions relative to the Control and High burn treatment across all time points (Figure 2c), resulting in higher cumulative N_2_O emissions in the Low burn treatment (3.11 ± 0.17 µg N-N_2_O g^-1^) than the Control (1.00 ± 0.98 µg N-N_2_O g^-1^) and High burn (0.88 ± 0.38 µg N-N_2_O g^-1^; Figure 2c,d; p < 0.03, LME). However, N_2_O emissions in the High burn treatment did not differ from those in the Control (p = 0.98, LME). Limited detection of the ^15^NH_4_^+^ tracer added to soils in the N_2_O we measured indicated that less than 1% of the N_2_O was directly produced by the oxidation of NH_4_^+^ (Table S2).

Net NH_4_^+^ production rates did not differ between the Low burn and the Control (p = 0.47, LME); however, the High burn stimulated net NH_4_^+^ consumption throughout the experiment (p < 0.001 High burn vs. Control; Figure 3a,b). Net NH_4_^+^ production was negatively correlated with peak burn temperatures (Figure S6a, p = 0.004, LME). In contrast to net NH_4_^+^ production rates, all treatments showed net NO_3_^-^ consumption; net nitrification rates in the Control, Low burn, and High burn all decreased from week one after burning until the end of the experiment (p < 0.0001, LME; Figure 3c). Overall, cumulative net nitrification (Figure 3d) was lowest in the Low burn relative to the High burn and the Control (p = 0.003, LME).

**Figure 3.**
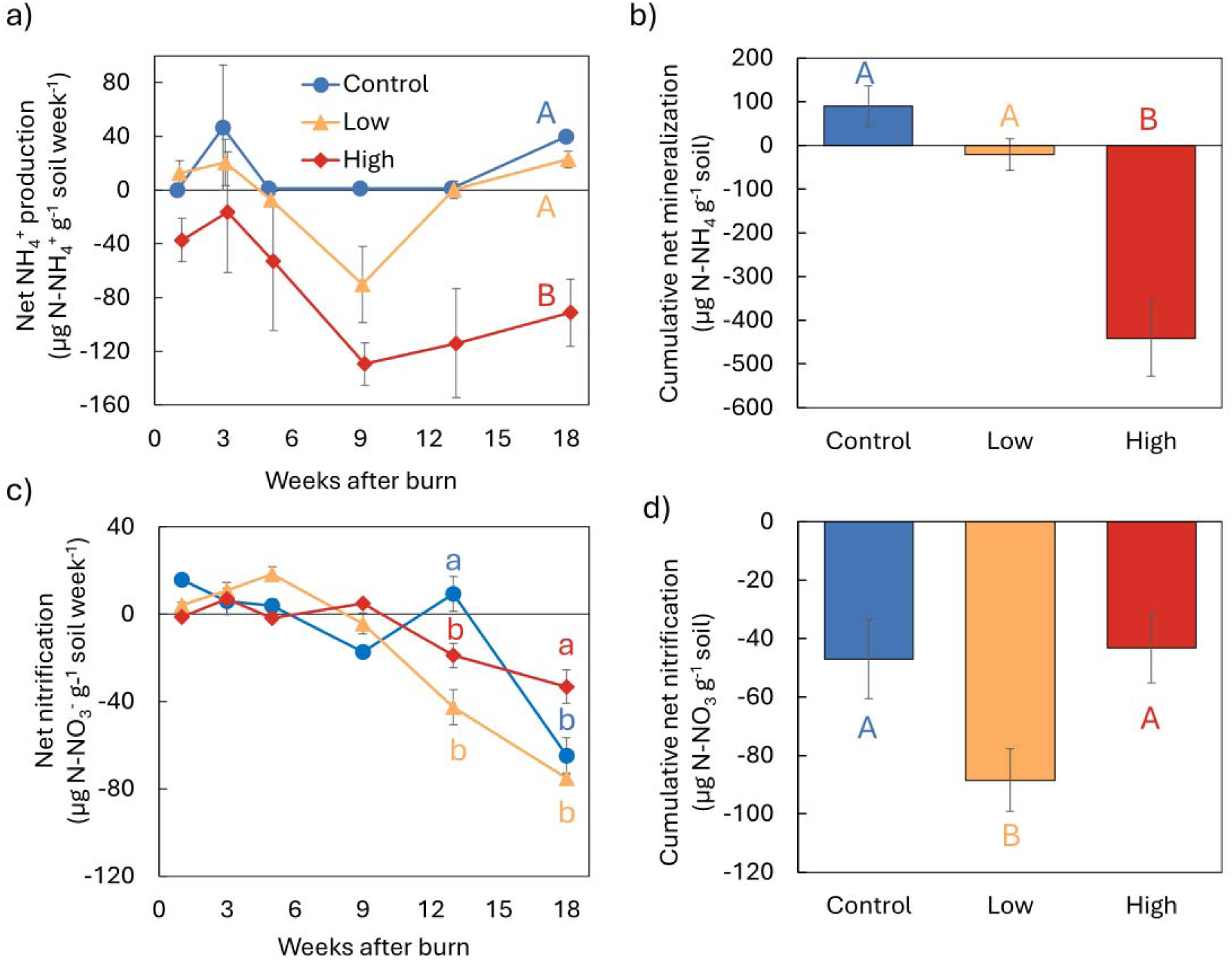
Net nitrogen transformation rates across burn intensity treatments over an 18-week incubation. Measured dynamics include time-series rates for (a) net NH_4_^+^ production and (c) net nitrification, alongside cumulative totals for (b) net NH_4_ ^+^ production and (d) net nitrification integrated over the 18 week incubation. Rates were calculated from KCl-extracted NH_4_^+^ and NO_3_^-^ concentrations measured before and after 7-day NO/N_2_O flux incubations shown in Figure 2. Pyrocosm treatments include unburned Control, Low burn intensity (peak temperature of 100 ± 20 °C at 5 cm depth), and High burn intensity (244 ± 7 °C at 5 cm depth). Different letters denote statistically significant differences (p < 0.05) based on linear mixed- effect models for time-series rates (a, c) and one-way ANOVAs for cumulative totals (b, d; lowercase letters indicate treatment differences within individual time points; uppercase letters represent main treatment effects). Overlapping points were jittered for error bar visibility. Error bars represent standard errors (n = 4).

### 3.3 Soil Carbon Pools

The High burn treatment significantly lowered C content relative to both the Control and Low burn (p < 0.03, LME; Figure 4a), whereas the Low burn had no effect on C content (p = 0.60, LME). In contrast to soil C content, the immediately respirable carbon (IRC) pool was greater in both the Low and High burns than in the Control during weeks 1 and 3 (p < 0.009, LME; Figure 4b), but the effect was short-lived and no longer detected for the remaining 18 weeks (p > 0.36). However, across all time points, the Low burn reached greater cumulative IRC than the Control (p = 0.034, LME), whereas the High burn had no effect on cumulative IRC (p = 0.13, LME; Figure 4c). Low intensity burns lowered microbial biomass C by 25% (p = 0.0082) and High intensity burns by 50% (p < 0.0001) relative to the Control one week after burning (Figure 4d), with burn effects on microbial biomass persisting for the 18 weeks of measurements. Microbial biomass C was also lower in the High burn treatment than in the Low burn (p = 0.0005, LME; Figure 4d).

**Figure 4.**
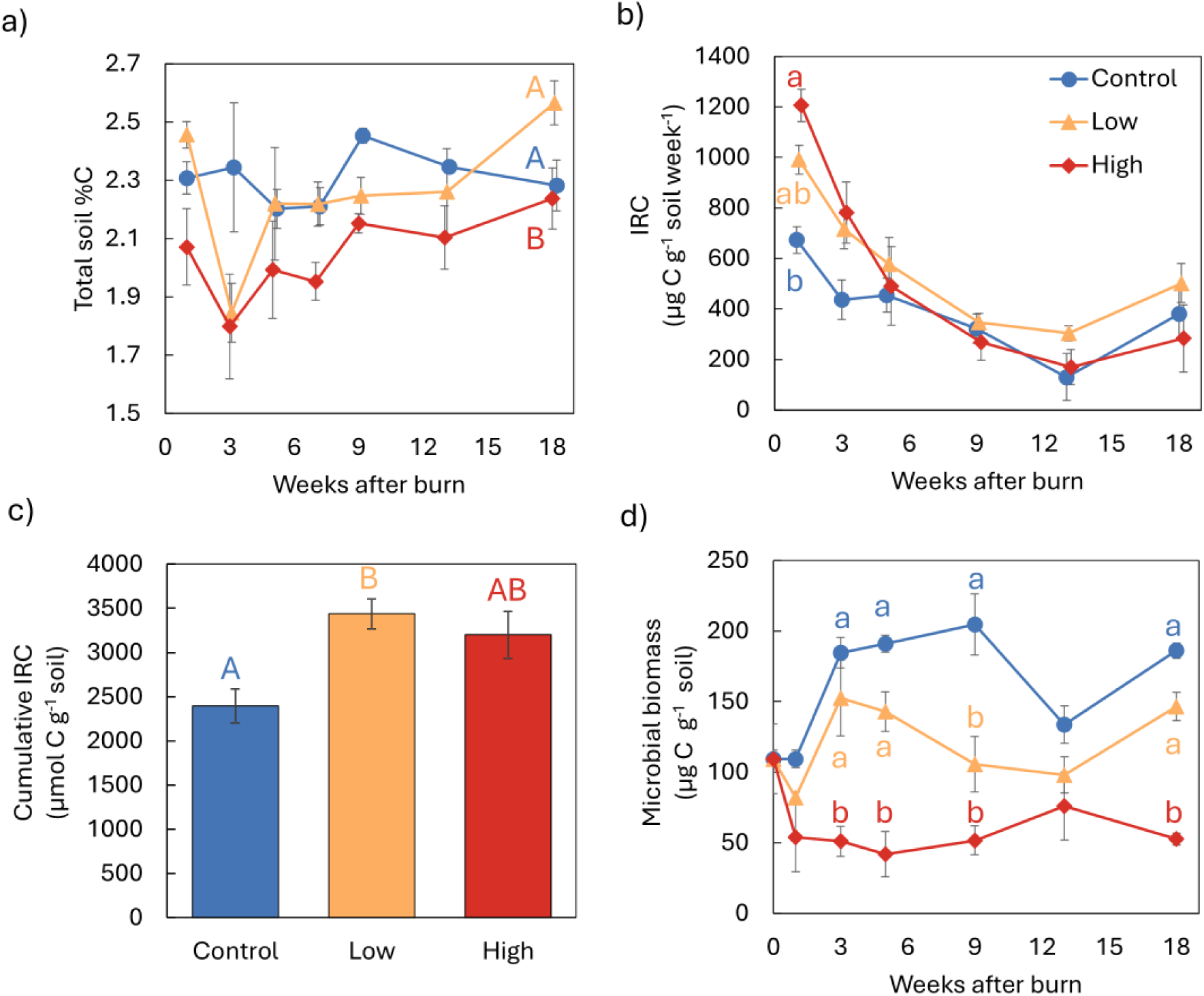
Soil carbon pools, respiration, and microbial biomass across burn intensity treatments over an 18-week incubation. Panels display (a) total soil % C, (b) immediately respirable C (IRC) derived from 7-day CO_2_ flux incubations, (c) cumulative IRC integrated over the 18 week incubation, and (d) microbial biomass C determined by chloroform fumigation. Pyrocosm treatments include unburned Control, Low burn intensity (peak temperature of 100 ± 20 °C at 5 cm depth), and High burn intensity (244 ± 7 °C at 5 cm depth). Different letters denote statistically significant differences (p < 0.05) based on linear mixed-effect models for time-series rates (a, b, d) and one-way ANOVAs for cumulative IRC (c; lowercase letters indicate treatment differences within individual time points; uppercase letters represent main treatment effects). Overlapping points were jittered for error bar visibility. Error bars represent standard errors (n = 4).

### 3.4 Extracellular Enzyme Activities

Potential hydrolytic enzyme activity generally decreased as burn intensity increased (Figure 5). Specifically, enzyme activity one week after burning in the Low burn was significantly lower than the Control: AG decreased 26% (p = 0.0008, using LMEs for all enzyme activities), BG decreased 33% (p = 0.0023), CBH decreased 63% (p = 0.006), NAG decreased 47% (p = 0.005), and PHO decreased 42% (p < 0.0001). Similarly, enzyme activity one week after burning in the High burn was significantly lower than both the Low burn and the Control: AG decreased 74% (p < 0.0001), BG decreased 87% (p < 0.0001), CBH decreased 98% (p < 0.0009), NAG decreased 85% (p < 0.0058), and PHO decreased 97% (p < 0.0001) relative to Control. After this initial burn effect, AG, BG, CBH, NAG, and PHO activities gradually increased over the 9 – 13 week period in the High burn but remained relatively constant in the Low burn and Control (Figure 5), but stabilized well below unburned levels after 18 weeks.

**Figure 5.**
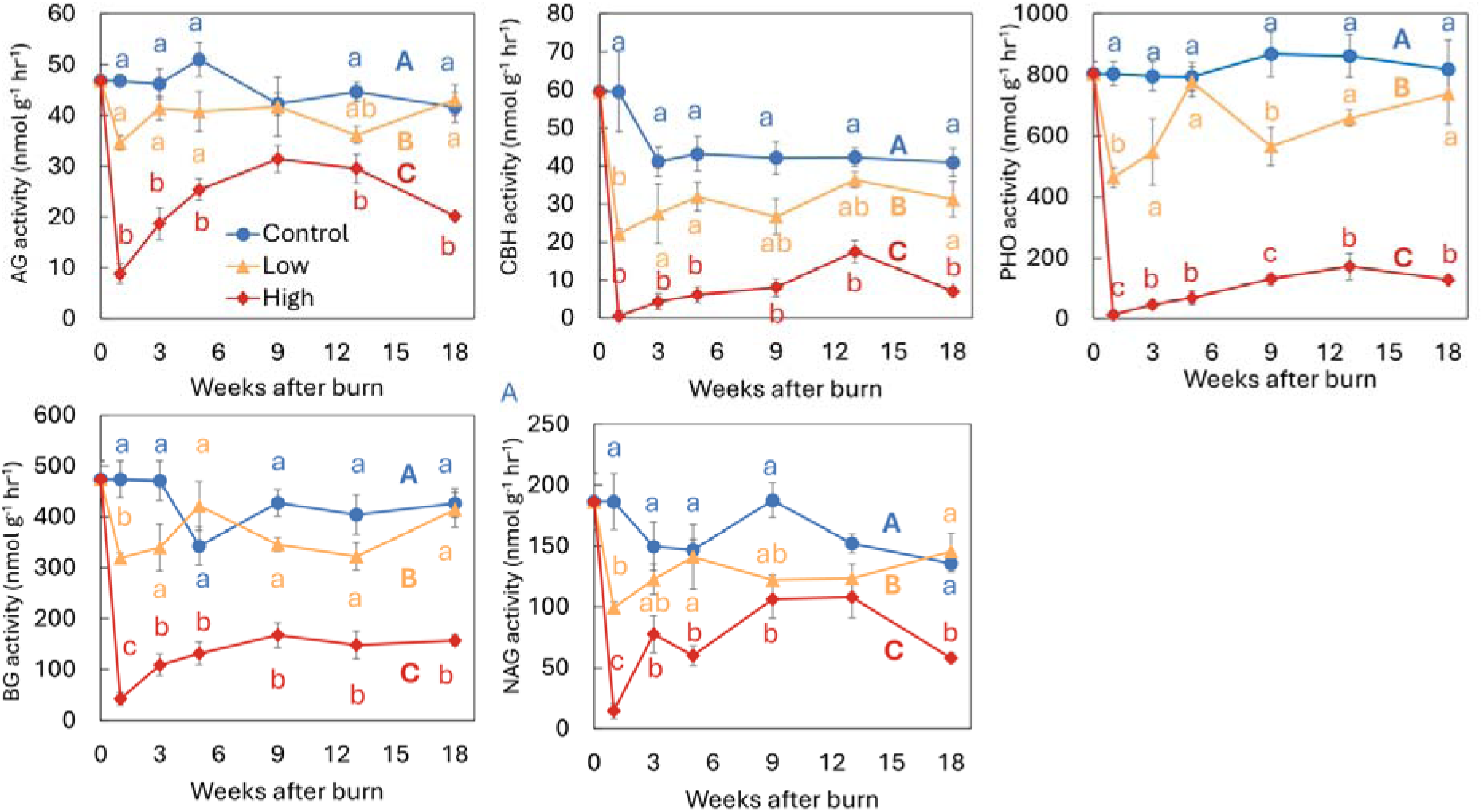
Potential extracellular hydrolytic enzyme activities across burn intensity treatments over an 18-week incubation. Target enzymes include C degrading enzymes α-glucosidase (AG), β-glucosidase (BG), and 1,4-β-cellobiohydrolase (CBH); the P acquisition enzyme phosphomonoesterase (PHO), and the N acquisition enzyme N-acetyl-β-d-glucosaminidase (NAG). Pyrocosm treatments include unburned Control, Low burn intensity (peak temperature of 100 ± 20 °C at 5 cm depth), and High burn intensity (244 ± 7 °C at 5 cm depth). Different letters denote statistically significant differences (p < 0.05) based on linear mixed-effect models for time-series rates (lowercase letters indicate treatment differences within individual time points; uppercase letters represent main treatment effects). Overlapping points were jittered for error bar visibility. Error bars represent standard errors (n = 4).

## 4. Discussion

Post-fire soils can persist as major sources of trace N gas emissions long after fires are extinguished, yet the drivers regulating post-fire N fluxes remain unresolved across studies (Stephens & Homyak, 2023). Using controlled pyrocosms to isolate burn intensity, we show that intensity exerts predictable, long-lasting effects on soil physicochemical and biological drivers— including soil pH, extractable NH_4_ ^+^ and NO_3_ ^-^, soil C availability (both total C and immediately respirable C; IRC), microbial biomass C (MBC), and potential extracellular enzyme activities— that persist for at least 18 weeks post-fire (Figures 6). Importantly, trace gas emissions showed divergent responses to burn intensity. While overall NO emissions were statistically unresponsive to burn intensity, mean fluxes trended higher in burned soils, and elevated soil moisture during flux incubations likely enhanced in-soil NO consumption and reduction to N_2_O, masking potential treatment effects. In contrast, N_2_O emissions showed a threshold response, increasing under Low burn intensity but declining under High burn intensity. The stimulation of N_2_O emissions at low intensity was driven by a combination of retained microbial biomass and enhanced substrate availability (i.e., IRC and NO_3_^-^), alongside negative net nitrification rates signaling active NO_3_ ^-^ consumption via denitrification. Beyond this low-intensity threshold, severe burns suppressed N_2_O emissions by depleting microbial biomass and substrate pools essential for denitrification (both C and N). Together, these results suggest that post-fire N_2_O emissions depend strongly on burn intensity up to a resource-limiting threshold governed by microbial and substrate limitation, whereas NO release is constrained by moisture-driven consumption that can override burn intensity effects during wet post-fire periods.

**Figure 6.**
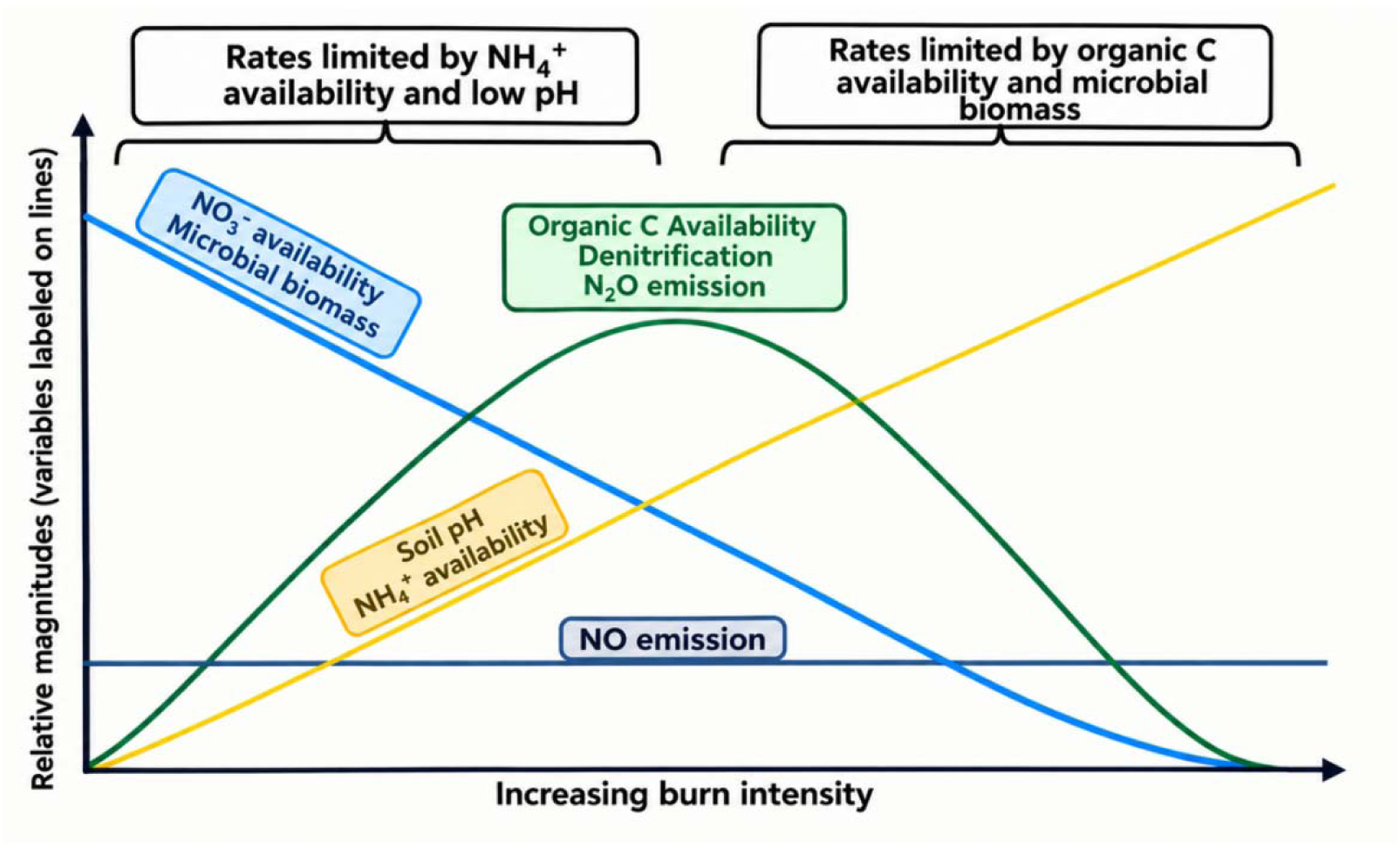
Conceptual summary of drivers shaping post-fire soil nitrogen (N) emissions across a burn intensity gradient. Process limitations on cycling rates are annotated above the plot. Nitric oxide (NO) emissions were not significantly affected by burn intensity despite fire-driven increases in soil pH and NH_4_^+^ availability. In contrast, nitrous oxide (N_2_O) emissions showed a non-linear threshold response, peaking under low burn intensity where soil pH, NO_3_^-^, and organic C availability were optimal. At higher burn intensities, denitrification and N_2_O emissions were constrained by NO_3_^-^ and labile C limitation, alongside reductions in microbial biomass.

Despite greater NH_4_^+^ availability to support nitrifiers (Figure 1c) and subsequent NO_3_^-^ accumulation in burned soils (Figure 1d), we did not detect significant increases in NO emission with burn intensity—a finding that contrasts with other grassland fire studies where post-burn NO fluxes increased markedly (Stephens & Homyak, 2023). However, although non-significant, mean NO emissions followed the expected pattern (i.e., High > Low > Control), and treatment differences may have started to emerge toward the end of the 18-week incubation (Figure 2a).

We attribute this lack of statistical significance to soil moisture differences between the “parent” pyrocosms and the laboratory flux incubation assays where we measured trace gases. Soil moisture in the pyrocosms was maintained below 50% WHC (Figure 1b), whereas flux incubations were initiated at 80% WHC and allowed to dry over 7 days (Figure S4b).

Consequently, while nitrification was active in the pyrocosm soils (Figure 1d), the high soil moisture during flux assays likely suppressed net NO evasion by stimulating NO consumption (e.g., reduction to N_2_O), inhibiting net nitrification as shown by negative net nitrification rates during incubations (Figure 3c). Thus, while burn intensity increased the NH_4_^+^ substrate pool to support nitrifiers, elevated soil moisture during flux assays likely facilitated in-soil NO consumption, masking treatment-driven emissions.

Several lines of evidence suggest that fire-induced shifts in soil pH and inorganic N availability stimulated nitrification and NO_3_^-^ availability, driving higher N_2_O emissions from denitrifiers. First, we note that our ^15^NH_4_^+^ tracer experiment confirmed that nitrifiers did not often produce more than 1% of the N_2_O directly (Table S2). Instead, nitrifiers, played an essential indirect role by generating the NO_3_^-^pool in the pyrocosms that denitrifiers consumed once soil moisture reached 80% WHC during flux incubations (Figures 1d), resulting in negative net nitrification rates (Figure 3c,d). As burn intensity increased, soil pH and NH_4_^+^ availability were positively related to peak burn temperatures (Figure S6c,d), consistent with well-documented post-fire dynamics (Ubeda et al. 2005; Fultz et al., 2016; Reinhart et al., 2016; Pereira et al., 2017; Cheng et al., 2025). Elevating soil pH shifts the equilibrium between NH_4_^+^ and NH_3_ towards NH_3_ (pK_a_ = 9.25, Krichels et al., 2023), the primary substrate oxidized by nitrifiers (Egan et al., 2018). Direct support for increased nitrification comes from the increase in net NH_4_^+^ consumption with burn intensity (Figure 3a,b) and the accumulation of extractable NO_3_^-^ after week 9 (Figure 1d). Because post-fire microbial biomass remained suppressed (Figure 4d; Zhilik et al., 2026), NH_4_^+^ immobilization was likely negligible, indicating that NH_4_^+^ consumption primarily reflected active nitrification. Ultimately, fire-enhanced nitrification increased the soil NO_3_ ^-^ pool, promoting denitrifier-derived N_2_O emissions that were also consistent with the late-stage temporal increase in NO emissions observed in the High burn treatment by week 18.

The non-linear response of N_2_O emissions as burn intensity increased likely reflects a threshold between processes stimulated by microbes and substrate limitation. To rule out differing oxic states as a factor influencing this non-linear response in N_2_O emissions, we note that WHC was kept consistent across our pyrocosms and flux measurements. This suggests the Low burn treatment generated conditions that favored denitrification: while microbial biomass was lower than the unburned Control, NO_3_^-^ availability was similar to the Control, with soil pH, total C, and IRC increasing due to the deposition of ash and partially combusted organic matter (Nelson et al., 2022; Dalias et al., 2024). In contrast, High burn intensity crossed a biological threshold, where the combustion of organic C, volatilization of NO_3_^-^through combustion, and decrease in microbial biomass constrained denitrification. Such stimulation of denitrification under low burn intensity—and its limitation under high burn intensity—was consistent with net nitrification dynamics, which served as a stronger net sink for NO_3_^-^ in the Low burn than in the Control or High burn treatments. We expect that the severe loss of microbial biomass decreased microbial NO_3_ ^-^ immobilization, thus we assume that NO_3_^-^ consumption reflects active denitrification (plants were not present in pyrocosms). Overall, this threshold response of denitrification to fire suggests that post-fire N_2_O emissions have a non-linear relationship to burn intensity, helping to explain why previous grassland soil field studies have observed divergent post-fire N_2_O responses ranging from substantial increases (Levine et al., 1988) to no observable change (Andersson et al., 2003; Castaldi et al., 2010; Livesley et al., 2011).

Potential extracellular enzyme activities further suggested that C and N limitation constrained denitrification following high-intensity burns. One week into our measurements, the potential activities of five hydrolytic enzymes decreased systematically with increasing burn intensity. Over the following weeks, however, enzyme activities in the High burn treatment rose significantly, whereas those in the Control and Low burn treatments remained relatively unchanged. Because soil microbes upregulate extracellular enzyme synthesis primarily to acquire scarce metabolic resources (Allison et al., 2005), this temporal increase suggests that High burn intensity severely restricted microbial access to labile organic C, compelling surviving microbes to invest in C-acquisition enzymes (Li et al., 2021). This is consistent with our observation that total C decreases in the High burn relative to the Control, but IRC in the High burn does not differ from the Control. Although enzymes like β-N-acetylglucosaminidase and phosphatase are conventionally classified as N- and P-acquiring, microbes frequently deploy them to harvest organic C (Mori et al., 2021; Lidbury et al., 2022). Overall, the elevated investment in C-acquiring enzymes further suggests that microbial respiration and denitrification were also constrained by labile C availability.

## 5. Conclusions

By isolating burn intensity under controlled conditions using a representative grassland soil, we show that post-fire N_2_O emissions follow a non-linear, fire intensity-dependent threshold. Low burn intensity stimulated a three-fold increase in N_2_O emissions, driven by fire-induced increases in soil pH, total C, immediately respirable C (IRC), and NO_3_^-^ availability that favored denitrification. Under high burn intensity, however, N_2_O emissions were constrained by lower microbial biomass, labile C, and NO_3_^-^ pools, offsetting the stimulatory effect of elevated pH (Figure 6). In contrast to N_2_O, soil NO emissions remained unaffected across burn intensities, likely because elevated soil moisture during flux incubations promoted in situ NO consumption and reduction to N_2_O, masking potential burn-driven emission signals. Overall, we show that in our controlled pyrocosms, post-fire soil N_2_O fluxes are governed by a balance between pH-driven microbial processes and severe resource limitation resulting from changes in fire intensity, offering a mechanistic perspective for the wide variability in post-fire N emissions reported across the literature (e.g., Stephens & Homyak, 2023). Incorporating burn intensity into biogeochemical models and spatial management frameworks (such as Burned Area Emergency Response classifications) could improve regional estimates of post-fire N gas losses and help refine predictions of wildfire impacts under changing wildfire regimes.

## Acknowledgements

This research was funded by the U.S. Department of Energy (Office of Biological and Environmental Research award #DE-SC0023127 to PMH, JBE, MJW, SDA, and SIG). We thank the Santa Rosa Ecological Plateau in Riverside County for site access, as well as University of California, Riverside (UCR) Agricultural Operations and Peggy Mauk for infrastructure and facility support. We are grateful to Krish Patel, Ryan Quaal, and Esbeiry Cordova-Ortiz for assistance with soil collection and pyrocosm burns, and to UCR undergraduate researchers Yareli Olazabal, Ciara Soto, Krish Patel, Ellen Nguyen, Rahma Ahmed, and Yanira Herrera for laboratory support with biogeochemical measurements.

Additionally, we thank Aral Greene and Jamie Irby for methodological training, Chloe Reid and Valerie Gonzalez for assistance with enzyme activity and elemental (%C) analyses, and Maggi Laan, Mason Laurin, Kai Huang, and Jingwen Xu for constructive feedback on the manuscript.

