## Supplemental Materials for "Burn intensity regulates post-fire effects on soil N_2_O emissions: A grassland soil pyrocosm study"

Supplemental Methods

Background δ^15^N was determined by averaging δ^15^N from Control and Low burn pyrocosm soil samples without adding ^15^NH_4_^+^ tracer (n = 15). The mass of N_2_O produced by nitrification was calculated by:

R_sample_ = (((δ^15^N/1000)+1) x R_standard_)

Where R_standard_ = 0.0036765

Atom % ^15^N-N_2_O = R_sample_/(R_sample_ +1) x 100

Atom % of tracer incorporated in N_2_O = (atom % of ^15^N-N_2_O in sample) – (atom % of ^15^N-N_2_O in background)

Mass N_2_O produced by nitrification = ((mass N_2_O produced)*(atom % of incorporated

tracer)/100)/(atom % tracer added to sample)

Deviation from background was determined by subtracting background δ^15^N (47.88) from all δ^15^N values. ^15^NH_4_ tracer was added to 2 atom % of KCl-extractable NH_4_^+^ measured in each sample, added once in the beginning of each incubation during initial wetting to 80% WHC. Pyrocosm treatments include unburned Control, Low burn intensity (peak temperature of 100 ± 20 ^o^C at 5 cm depth), and High burn intensity (244 ± 7 ^o^C at 5 cm depth).

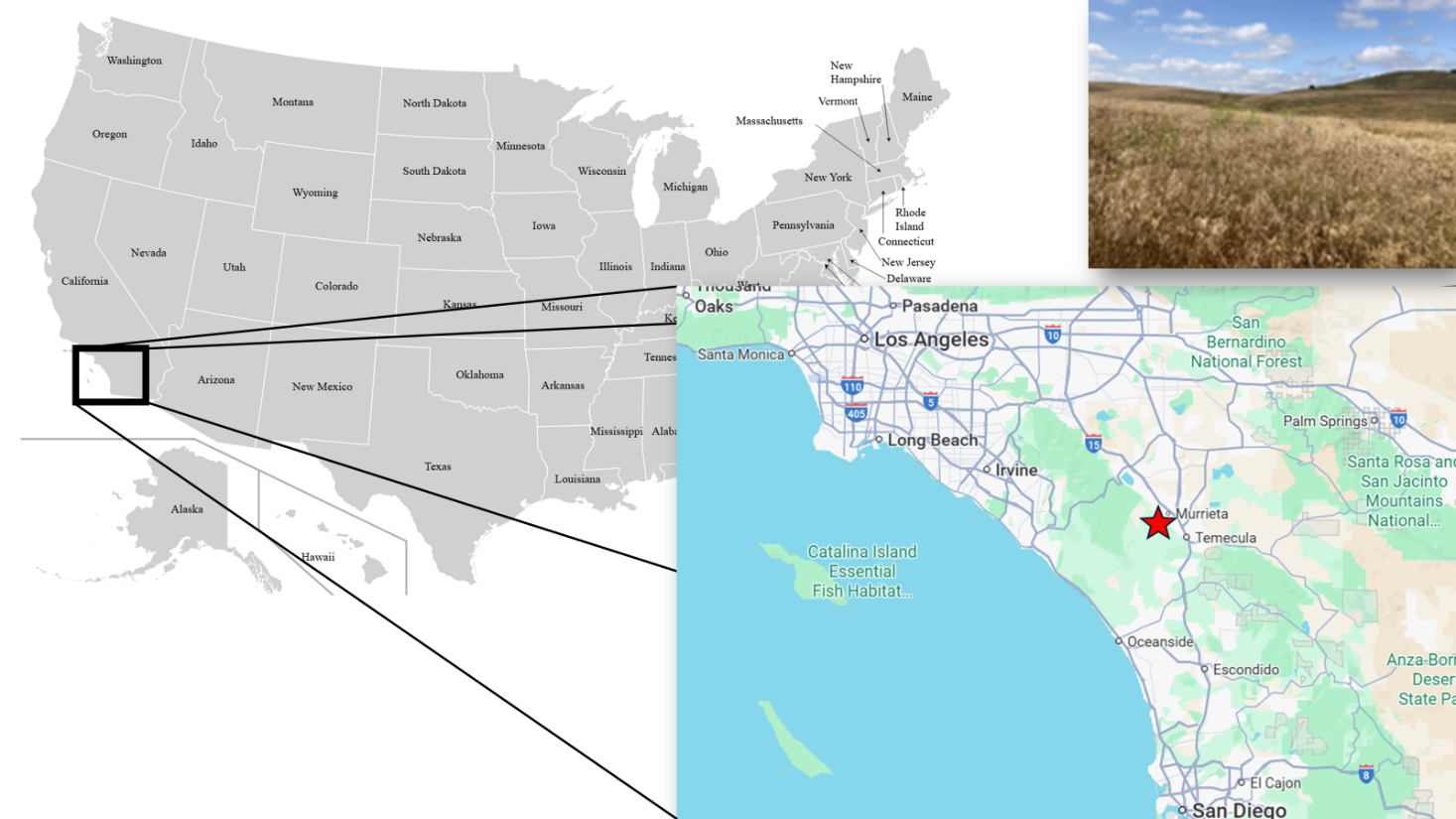

**Figure S1.** Map of Southern California, USA, showing the location of the soil collection site, Santa Rosa Plateau Ecological Reserve, and photo of the site. Soil was collected from 33°32'02.0"N 117°15'15.9"W (map image from Google Maps).

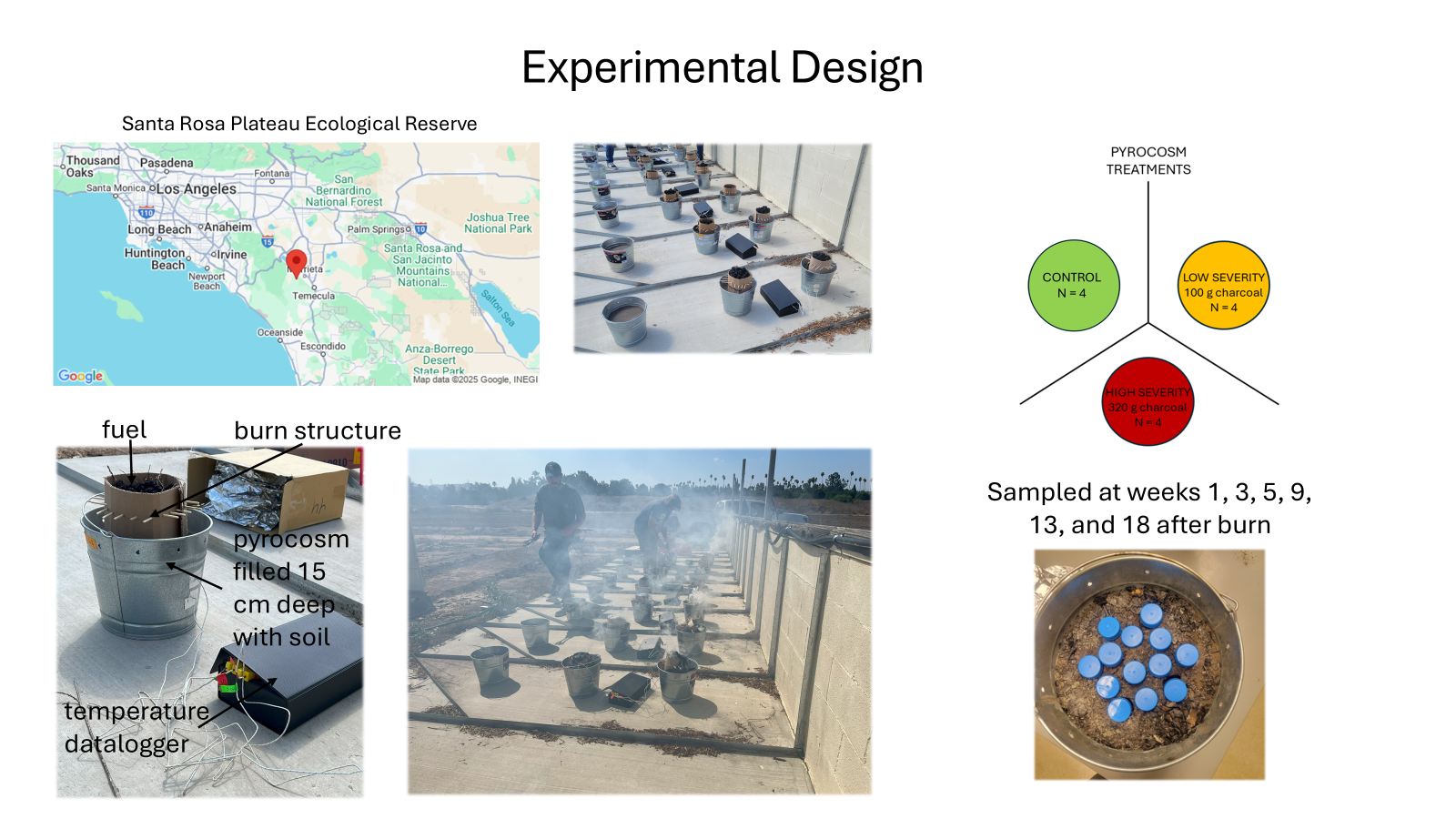

**Figure S2.** Pyrocosm setup showing the cardboard and wood structure used to contain the charcoal and control temperature profiles of both Low and High intensity burns. Thermocouples were installed at 5 and 10 cm depths.

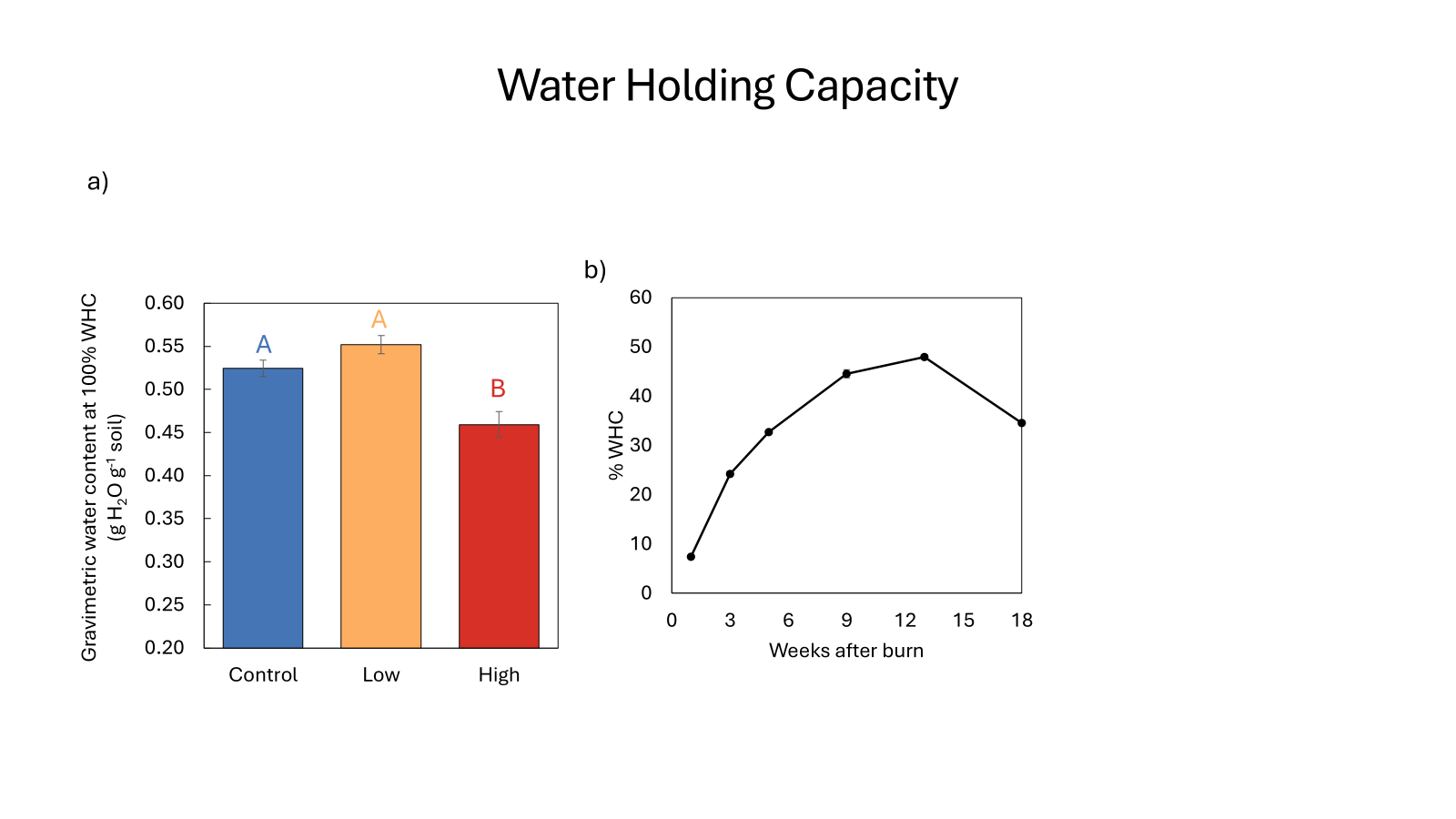

**Figure S3.** **Soil gravimetric water content at 100% water-holding capacity (WHC) one week post-fire**. Pyrocosm treatments include unburned Control, Low burn intensity (peak temperature of 100 ± 20 ^o^C at 5 cm depth), and High burn intensity (244 ± 7 ^o^C at 5 cm depth). Different letters denote statistically significant differences (p < 0.05). Error bars represent standard errors (n = 4).

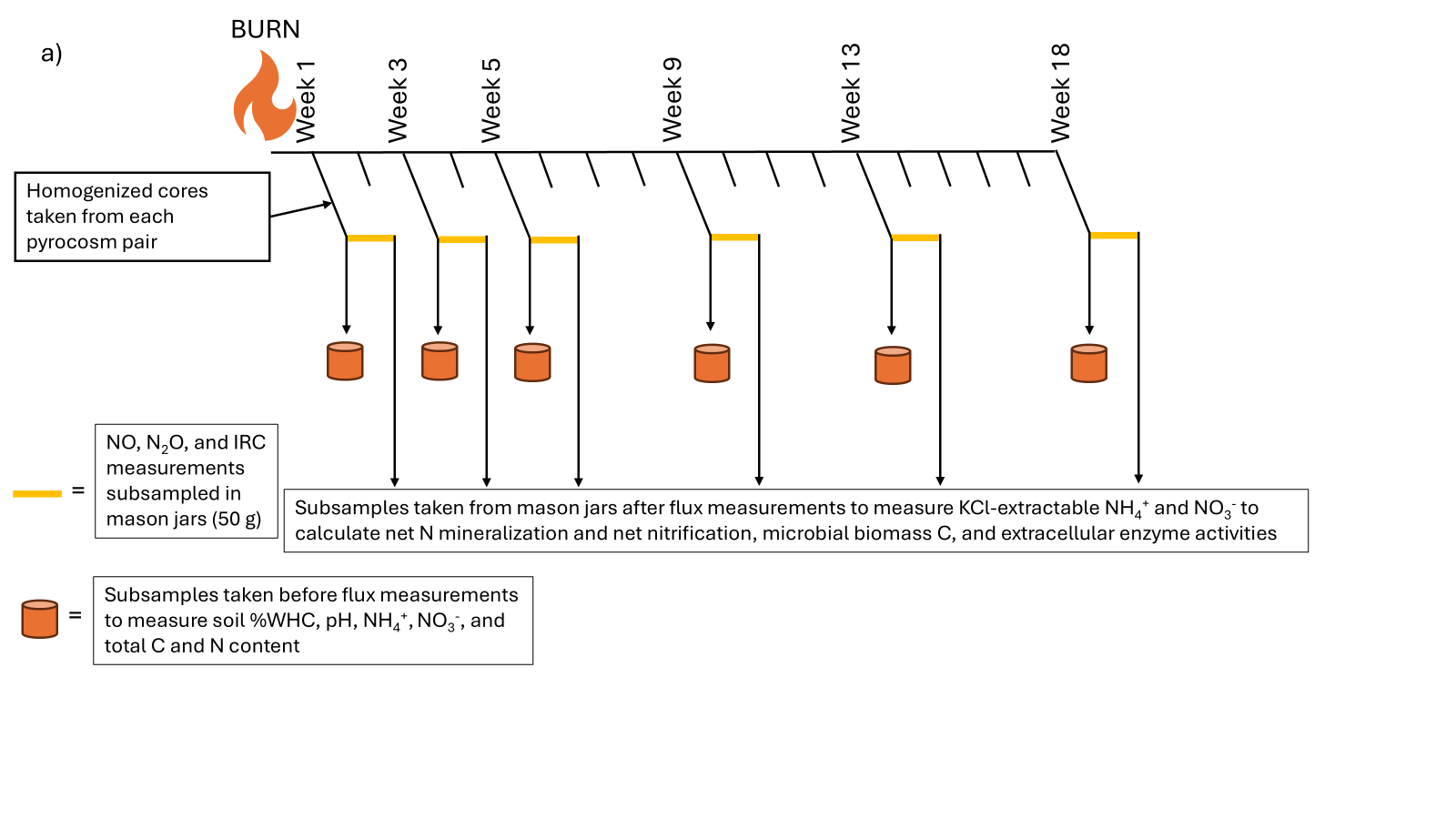

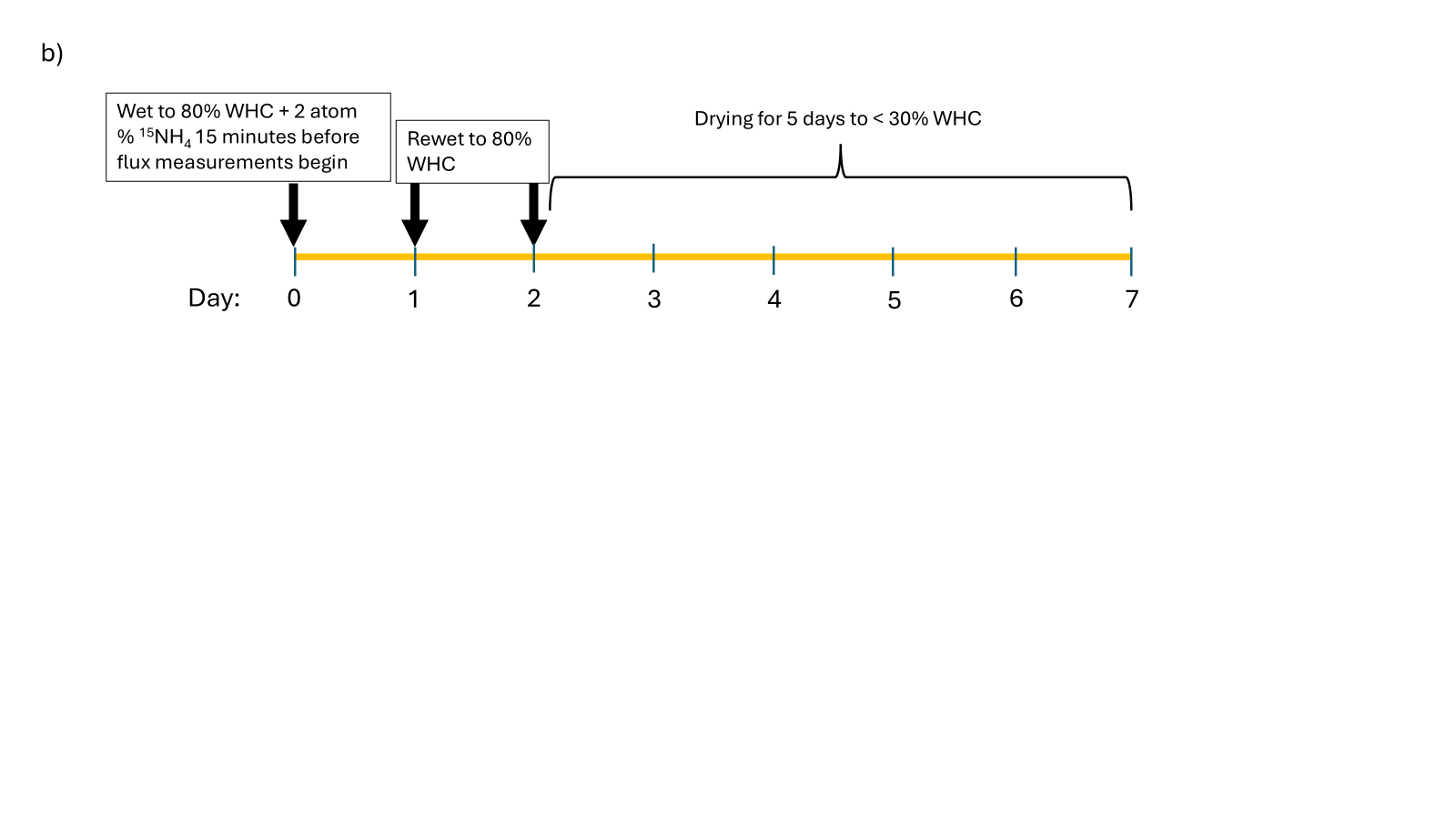

**Figure S4. Experimental timeline and sampling schematic for the 18-week pyrocosm incubation.** (a) Overview of the 18 week experimental schedule illustrating sampling timepoints, trace gas flux measurement windows (yellow lines), and soil subsampling for biogeochemical analyses conducted before (brown cylinders) and after 7-day flux incubations. (b) Zoomed-in detail of an individual 7-day trace gas flux measurement cycle corresponding to the incubation intervals in panel a.

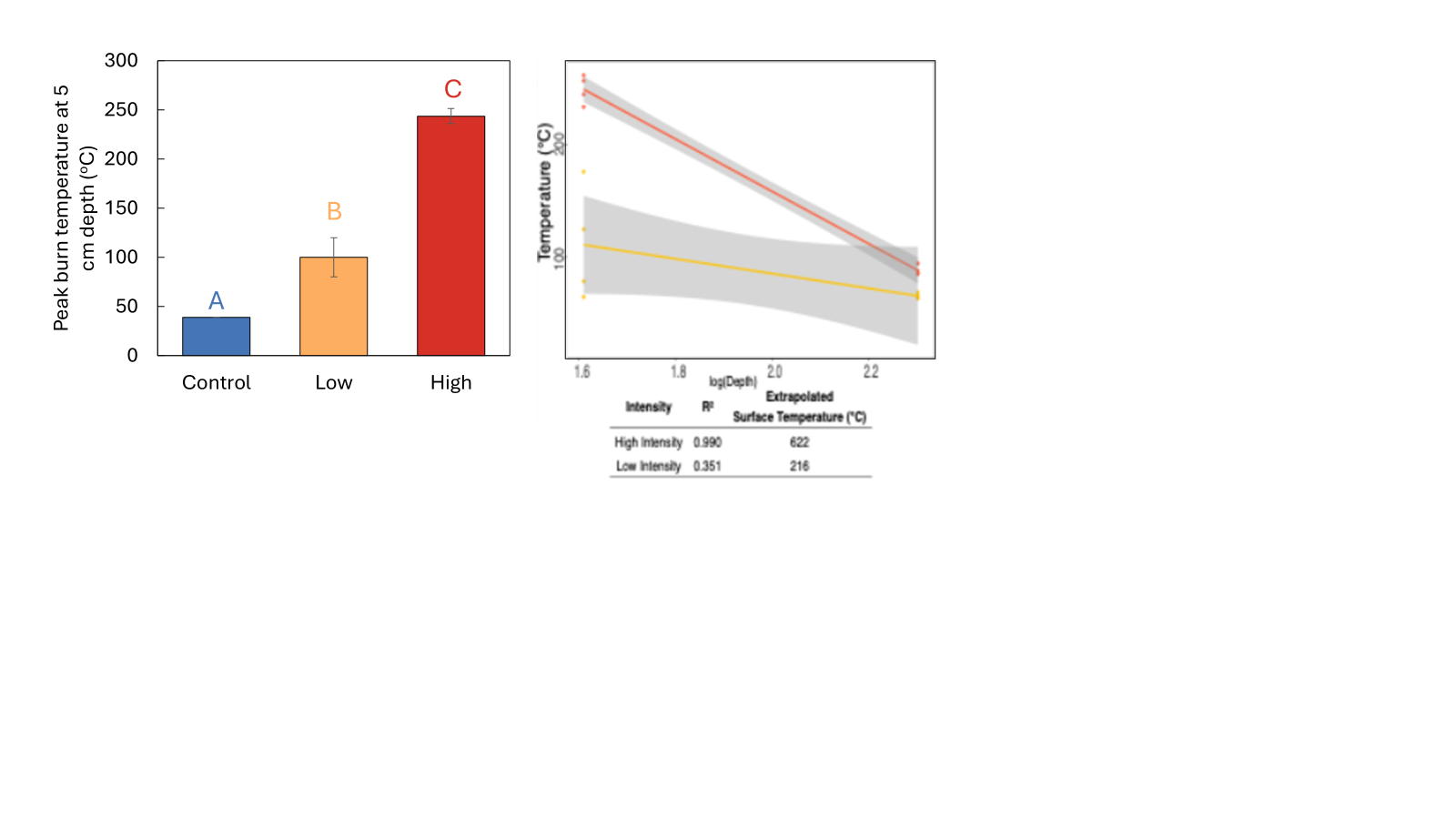

**Figure S5.** **Peak soil temperatures recorded at 5 cm depth across burn treatments and linear extrapolation to surface temperatures from 5 and 10 cm depth soil surface temperatures**. For unburned Control pyrocosms, the bar represents peak ambient air temperature on the day of burning. Peak subsurface temperatures differed significantly among all three treatments (p < 0.017). Different letters above bars denote statistically significant differences (p < 0.05) determined using a one-way ANOVA with post-hoc comparisons. Error bars represent standard errors (n = 8).

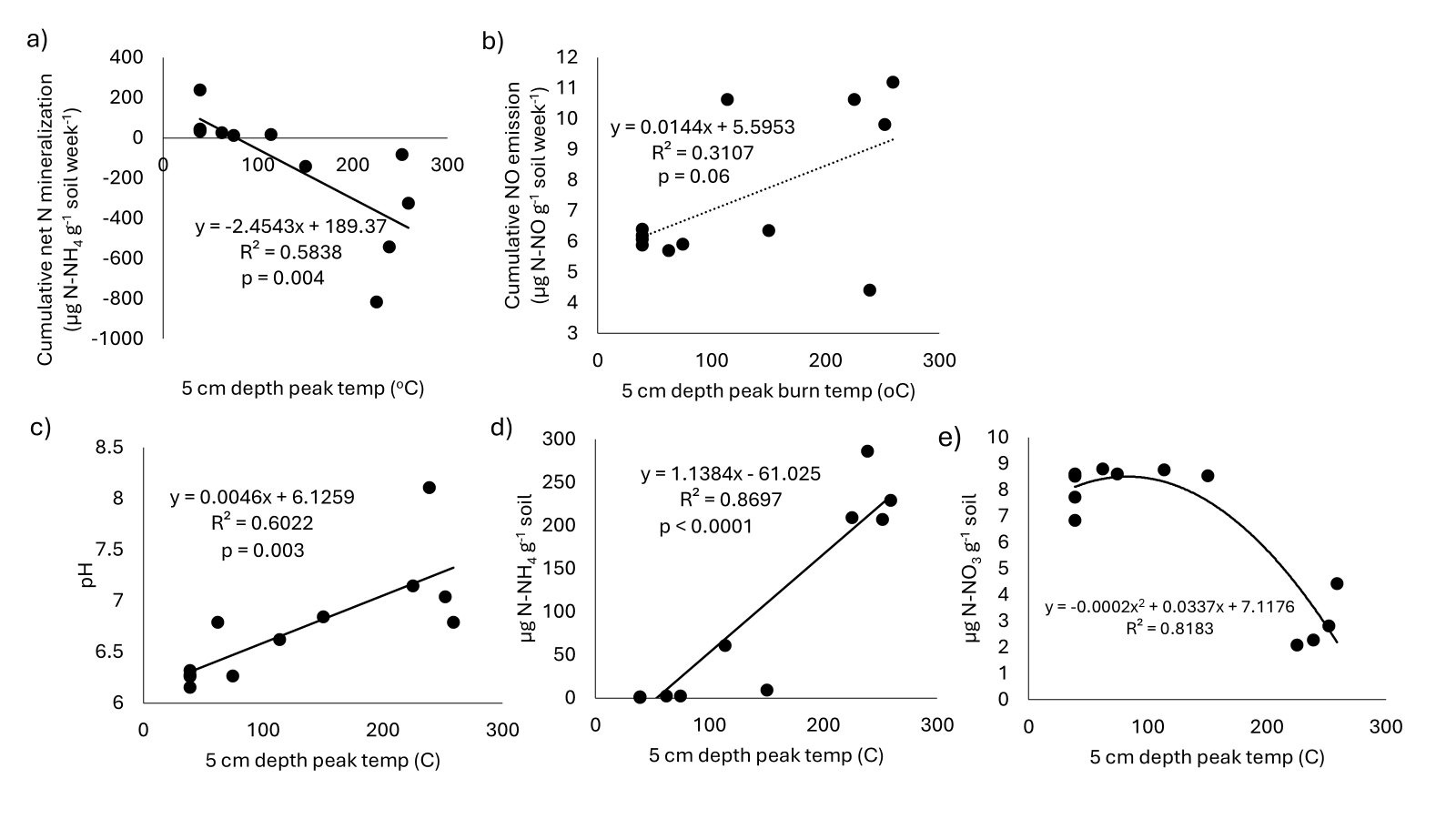
 **Figure S6.** **Linear and polynomial regressions between peak subsurface burn temperature (5 cm depth) and key soil biogeochemical properties**. Dependent variables include (a) cumulative net N mineralization, (b) NO emissions, (c) soil pH one week after burning, (d) extractable soil NH_4_^+^ one week after burning, and (e) extractable NO_3_^-^ one week after burning. For unburned Control pyrocosms, peak ambient air temperature on the day of burning is shown.

**Table S1.** Results of repeated measures ANOVAs and linear mixed-effect models testing for differences between treatments and time points in pyrocosm measurements. Asterisks indicate significance (p < 0.05).

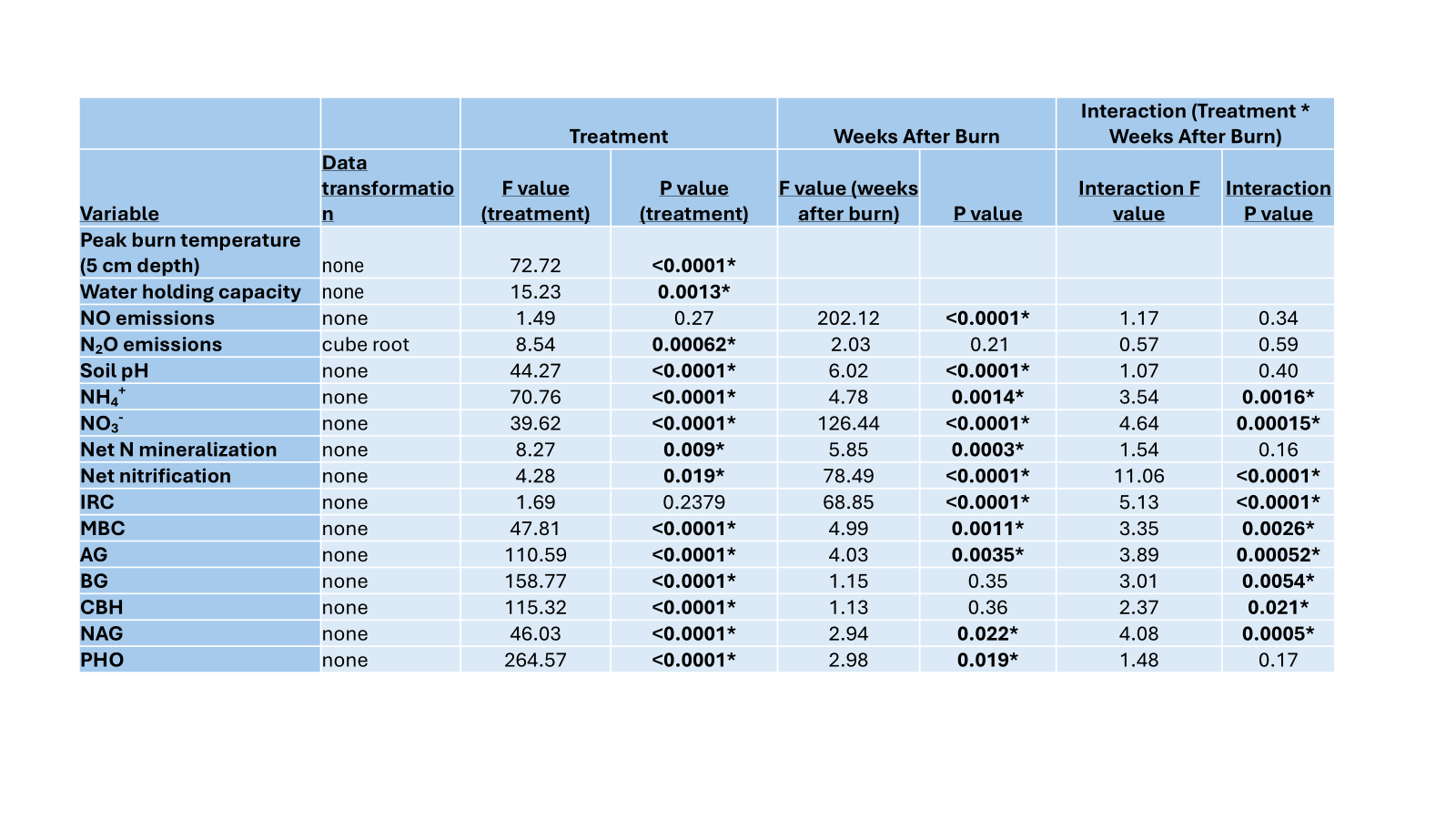

**Table S2.** Percentage of N_2_O produced by nitrification as determined by incorporation of ^15^NH_4_^+^ tracer in the N_2_O emitted from pyrocosms across burn intensity treatments over an 18-week incubation.

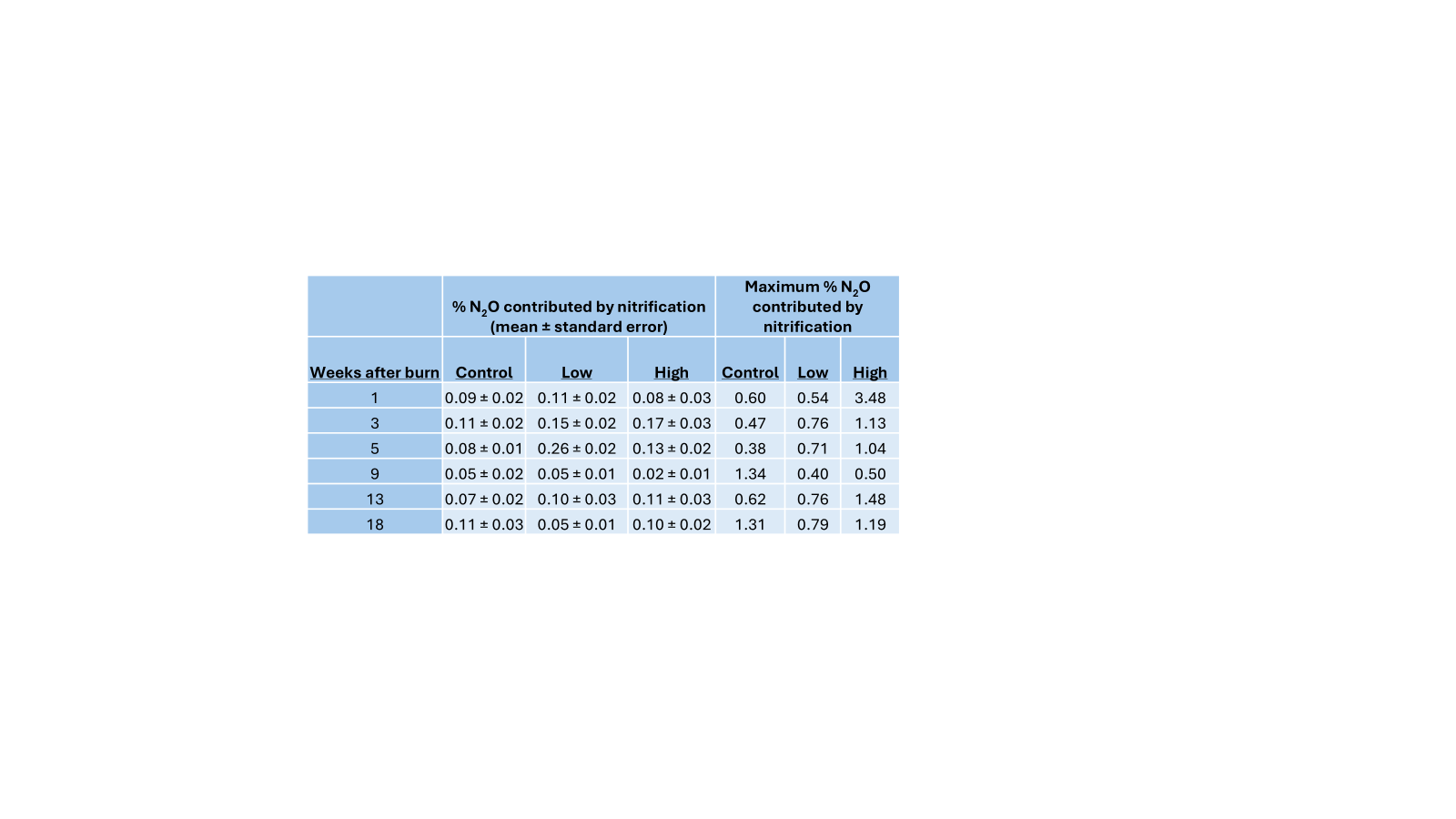
